# Genetic and transgenic reagents for *Drosophila simulans, D. mauritiana, D. yakuba, D. santomea* and *D. virilis*

**DOI:** 10.1101/096644

**Authors:** David L. Stern, Justin Crocker, Yun Ding, Nicolas Frankel, Gretchen Kappes, Elizabeth Kim, Ryan Kuzmickas, Andrew Lemire, Joshua D. Mast, Serge Picard

## Abstract

Species of the *Drosophila melanogaster* species subgroup, including the species *D. simulans, D. mauritiana, D. yakuba*, and *D. santomea*, have long served as model systems for studying evolution. Studies in these species have been limited, however, by a paucity of genetic and transgenic reagents. Here we describe a collection of transgenic and genetic strains generated to facilitate genetic studies within and between these species. We have generated many strains of each species containing mapped *piggyBac* transposons including an *enhanced yellow fluorescent protein* gene expressed in the eyes and a *phiC31 attP* site-specific integration site. We have tested a subset of these lines for integration efficiency and reporter gene expression levels. We have also generated a smaller collection of other lines expressing other genetically encoded fluorescent molecules in the eyes and a number of other transgenic reagents that will be useful for functional studies in these species. In addition, we have mapped the insertion locations of 58 transposable elements in *D. virilis* that will be useful for genetic mapping studies.

---

Ever since Alfred Sturtevant discovered *Drosophila simulans*, the sister species to *D. melanogaster*, in 1919, species of the *Drosophila melanogaster* species subgroup have played a central role in studies of evolution and speciation (Powell 1997; Barbash 2010). Most species of the subgroup display superficially similar anatomy, although all species can be distinguished by both qualitative and quantitative anatomical differences (Orgogozo and Stern 2009). In addition, the species display enormous variation in ecology and behavior, with some having evolved into ecological specialists on unusual food sources (R’Kha *et al.* 1991; Yassin *et al.* 2016).

One of the major advantages of this subgroup for evolutionary studies is that many of the species can be crossed to *D. melanogaster* to generate sterile hybrids and some can be crossed to each other to generate fertile hybrid females (Powell 1997). An unusual and important feature of these fertile pairs is that strains of each species can be found that share synteny across all chromosomes (Lemeunier and Ashburner 1976; Moehring *et al.* 2006a). This allows comprehensive genetic interrogation of the entire genome through recombination mapping. This is an uncommon feature for fertile pairs of *Drosophila* species; most species that have been examined exhibit major chromosomal inversions that are fixed between species (Powell 1997).

The combination of relatively straightforward genetics with diversity in anatomy, physiology and behavior has encouraged many groups to perform genetic analyses of these species (e.g. Liu *et al.* 1996; True *et al.* 1997; Macdonald and Goldstein 1999; Gleason and Ritchie 2004; Moehring *et al.* 2004, 2006a; b; Carbone *et al.* 2005; Gleason *et al.* 2005; Orgogozo *et al.* 2006; Cande *et al.* 2012; Arif *et al.* 2013; Peluffo *et al.* 2015). In the vast majority of cases, however, these studies have stopped after quantitative trait locus (QTL) mapping of traits of interest. One factor that has limited further genetic study of these traits is a limited set of genetic markers, which can facilitate fine-scale mapping. John True and Cathy Laurie established a large collection of strains carrying *P-element* transposons marked with a w^+^ mini-gene in a w-background of *D. mauritiana* (True *et al.* 1996a; b). These have been used for introgression studies (True *et al.* 1996b; Coyne and Charlesworth 1997; Tao *et al.* 2003a; b; Masly and Presgraves 2007; Masly *et al.* 2011; Arif *et al.* 2013; Tanaka *et al.* 2015; Tang and Presgraves 2015) and for high-resolution mapping studies (McGregor *et al.* 2007; Araripe *et al.* 2010), demonstrating the utility of dominant genetic markers for evolutionary studies. One limitation of these strains is that the w^+^ marker is known to induce behavioral artifacts (Zhang and Odenwald 1995; Campbell and Nash 2001; Xiao and Robertson 2016). We have also observed that mutations in the *white* gene and some w^+^ rescue constructs cause males to generate abnormal courtship song (unpublished data). Other pigmentation genes that are commonly used in *D. melanogaster* are also known to disrupt normal behavior (Bastock 1956; Kyriacou et al. 1978; Drapeau et al. 2006; Suh and Jackson 2007). It would be preferable, therefore, to employ dominant genetic markers that do not interfere with normal eye color or pigmentation.

We were motivated by the phenotypic variability and genetic accessibility of these species to establish a set of reagents that would allow, simultaneously, a platform for site-specific transgenesis (Groth *et al.* 2004) and reagents useful for genetic mapping studies. We therefore set out to establish a collection of strains carrying transposable elements marked with innocuous dominant markers for four of the most commonly studied species of the *D. melanogaster* species subgroup: *D. simulans, D. mauritiana, D. yakuba* and *D. santomea*. We chose the *piggyBac* transposable element to minimize bias of insertion sites relative to gene start sites (Thibault *et al.* 2004) and integrated transposable elements carrying *enhanced yellow fluorescent protein* (*EYFP*) and *DsRed* driven by a *3XP3* enhancer that is designed to drive expression in the eyes (Horn *et al.* 2003). A large subset of the lines described here also include a phiC31 *attP* landing site to facilitate site-specific transgene integration. Here we describe the establishment and mapping of many lines of each species carrying *pBac{3XP3::EYFP,attP}* and *pBac{3XP3::DsRed}* (Horn *et al.* 2003). We have characterized a subset of the *pBac{3XP3::EYFP,attP}* lines from each species for phiC31 integration efficiency of plasmids containing an *attB* sequence. In addition, we have integrated transgenes carrying the *even-skipped* stripe 2 enhancer to characterize embryonic expression generated by a subset of *attP* landing sites. We have employed CRISPR/Cas9 to knock out the *3XP3::EYFP* gene in a subset of lines to facilitate integration of reagents for neurogenetics. We also describe several other genetic and transgenic reagents that may be useful to the community, including the map positions for *pBac* transposons integrated in the *D. virilis* genome.

## Methods

**Transposable elements employed:** We used *piggyBac* transposable elements (Horn *et al.* 2003) to mobilize markers to random locations within the genomes of *D. simulans white*[501] (San Diego Species Stock Center stock number 14021-0251.011), *D. simulans yellow*[1] *white*[1] (San Diego Species Stock Center stock number 14021-0251.013), *D. mauritiana white*^-^ (San Diego Species Stock Center stock number 14021-0241.60), *D. yakuba white*^-^ (San Diego Species Stock Center stock number 14021-0261.02), *D. santomea* STO CAGO 1482 (provided by Peter Andolfatto), and *D. virilis w*[50112] (San Diego Species Stock Center number 15010-1051.53). We constructed *pBac{3XP3::EYFP-attP}* by cloning a BglII fragment containing the *attP* site from *pM{3XP3-RFPattP’}* (Bischof *et al.* 2007) into the single BglII site of *pBac{3XP3::EYFPafm}* (Horn and Wimmer 2000).

We constructed *pBac* plasmids carrying a source of *P*-element transposase marked with *3XP3::EYFP* or *3XP3::DsRed* as follows. We digested the plasmid pACNNTNPII-S129A (Beall *et al.* 2002) with EcoRI and NotI and cloned the ~5kb fragment resulting from digestion into pSLFa1180fa (Horn and Wimmer 2000). This plasmid was digested with AscI or FseI and the ~5kb fragment was cloned into the AscI or FseI restriction sites of *pBac{3XP3::DsRed}* or *pBac{3XP3::EGFP,attP}* (Horn and Wimmer 2000) to generate *pBac{Pactin::Ptrsps, 3XP3::DsRed}* and *pBac{Pactin::Ptrsps 3XP3::EGFP,attP}*, respectively. These plasmids were injected into strains of *D. simulans* and *D. mauritiana*. We also injected *pBac{3XP3::DsRed}*(Horn *et al.* 2003) into strains of *D. simulans, D. mauritiana, D. yakuba*, and *D. santomea*. The complete sequences of *pBac{3XP3::EYFP-attP}*, *pBac{3XP3::DsRed}* and *phsp-pBac* are provided as Supplementary Material. These plasmids were co-injected with 250 ng/uL *phsp-pBac* (Handler and Harrell 1999), a heat-shock inducible source of *piggyBac* transposase, and one hour after injection embryos were heat shocked at 37°C for one hour. All embryo injections were performed by Rainbow Transgenic Flies Inc. G0 flies were backcrossed to un-injected flies of the same strain and G1 flies were screened for fluorescence in their eyes.

Fluorescence could be detected easily in the compound eyes and ommatidia in all of the *white*^-^ strains (*D. simulans, D. mauritiana, D. yakuba*, and *D. virilis*) using any dissecting microscope we tried with epi-fluorescence capability (Figure 1a). In flies with wild-type eye coloration, fluorescence in the compound eye is limited to a small spot of about ten ommatidia (Figure 1b). However, we found that fluorescence was very weak, and usually unobservable, in the eyes of flies with wild-type eye coloration using a Leica 165 FC stereomicroscope. This microscope uses “TripleBeam Technology” to deliver excitation light along a separate light path from the emission light. Unfortunately, the excitation light in this system appears to illuminate ommatidia adjacent to the ommatidia that are viewed for the emission light. Fluorescence can still be detected in the ocelli of these flies with this microscope, although this requires a bit more patience than when using a standard epi-fluorescence microscope to screen for fluorescence in the compound eyes.

**Figure 1.**
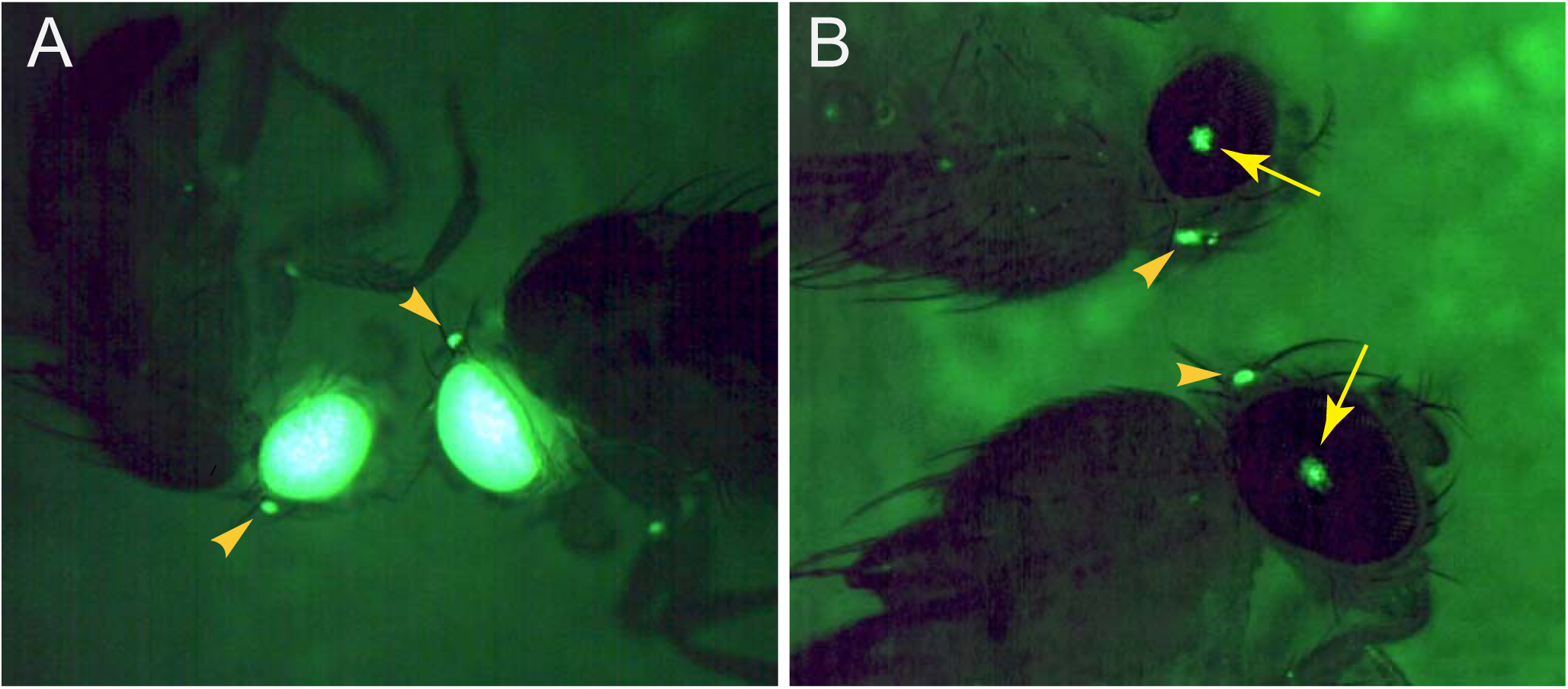
Appearance of EYFP fluorescence in fly eyes. (A) In flies carrying a *w*^-^ mutation, fluorescence is often intense and observable throughout the compound eye and in the (arrowheads). (B) In flies carrying wild-type eye coloration, fluorescence is observed in the compound eye as small dots including about 10 ommatidia (arrows) and in the ocelli (arrowheads).

**Mapping of transposable element insertion sites:** We mapped the genomic insertion sites of all *pBac* elements using both inverse PCR (Ochman *et al.* 1988) and TagMap (Stern 2016). Inverse PCR (iPCR) was not ideal for our project for several reasons. First, many isolated strains appeared to contain multiple insertion events, even though they were isolated from single G0 animals. These multiple events could sometimes be detected by segregation of offspring with multiple strengths of fluorescence in the eyes. In these cases, sometimes iPCR produced uninterpretable sequences and sometimes apparently only a single insertion event amplified. Second, many iPCR sequences were too short to allow unambiguous mapping to the genome. Third, sometimes iPCR reactions failed for no obvious reason. For all of these reasons, it was difficult to unambiguously map all of the *pBac* insertions with iPCR. We therefore developed and applied TagMap (Stern 2016) to map the insertion positions of all *pBac* elements. TagMap combines genome fragmentation and tagging using Tn5 transposase with a selective PCR to amplify sequences flanking a region of interest. This method provides high-throughput, accurate mapping of transposon insertions. Tagmap provided transposon insertion positions for all but a few strains. Transposable element insertion sites in the *D. simulans* and *D. mauritiana* strains were mapped to *D. simulans* genome release 2 (Hu *et al.* 2013), available from http://ftp.flybase.net/genomes/Drosophila_simulans/dsim_r2.01_FB2015_01/. Insertion sites in *D. yakuba* and *D. santomea* were mapped to *D. yakuba* genome release 1.3 (Clark *et al.* 2007), available from http://ftp.flybase.net/genomes/Drosophila_yakuba/dyak_r1.3_FB2014_03/. The actual genomes used for mapping and the mapped positions of the transposable elements are provided in the Geneious files supplied as Supplementary Files.

**Table 1.** Number of *attP* strains of each of five species that did not or did support integration of *attB* plasmids. Details are available in Supplementary File: Strains and Integration Efficiencies.xlsx.

| Species | Number of strains with zero integrants | Number of strains with at least one integrant |
| --- | --- | --- |
| *D. mauritiana* | 14 | 21 |
| *D. simulans* | 13 | 29 |
| *D. santomea* | 1 | 8 |
| *D. yakuba* | 1 | 19 |
| *D. virilis* | 9 | 0 |

**Mapping *pBac* transposon insertion sites in *D. virilis*:** We previously generated multiple *pBac*(*enhancer-lacZ*) insertions into *D. virilis* to study the *svb* gene (Frankel *et al.* 2012). However, none of these *pBac* (*enhancer-lacZ*) insertions have been mapped previously. These reagents may be useful for genetic mapping studies. We have therefore mapped positions of these inserts using TagMap. The larger scaffolds from the *D. virilis* CAF1 assembly project (http://insects.eugenes.org/species/data/dvir/) (Clark *et al.* 2007) have been mapped to Muller elements (Schaeffer *et al.* 2008). We combined this information with genetic linkage data to assemble approximately 159 Mbp of the *D. virilis* genome into the six Muller arms (unpublished data). We mapped insertion sites to this unpublished version of the *D.virilis* genome.

**Generation of a *D. santomea white-* allele:** We began to generate this collection of reagents prior to the availability of a *white*^-^ strain of *D. santomea.* However, soon after CRISPR/Cas9-mediated genome editing became available, we generated a *white*^-^ strain derived from *D. santomea* STO-CAGO 1482 as follows. *In vitro* transcribed Cas9 mRNA, generated with an EcoRI digested T7-*Cas9* template plasmid and the mMESSAGE mMACHINE T7 Transcription Kit (ThermoFisher Scientific), together with two gRNAs targeting the third exon of the *white* gene were injected into pre-blastoderm embryos by Rainbow Transgenics. The sequence for the T7-*Cas9* plasmid is provided as Supplementary Data. The gRNAs were generated by separate *in vitro* transcription reactions, using the MEGAscript T7 Transcription Kit (ThermoFisher Scientific), of PCR amplified products of the following forward and reverse primers: Forward primer CRISPRF-san-w12, 5’ GAA ATT AAT ACG ACT CAC TAT AGG CAA CCT GTA GAC GCC AGT TTT AGA GCT AGA AAT AGC; Forward primer CRISPRF-san-w17, 5’ GAA ATT AAT ACG ACT CAC TAT AGG GCC ACG CGC TGC CGA TGT TTT AGA GCT AGA AAT AGC; Reverse primer gRNA-scaffold, 5’ AAA AGC ACC GAC TCG GTG CCA CTT TTT CAA GTT GAT AAC GGA CTA GCC TTA TTT TAA CTT GCT ATT TCT AGC TCT AAA AC. All PCR reactions described in this paper were performed using Phusion High Fidelity DNA Polymerase (NEB) using standard conditions. Injected G0 flies were brother-sister mated and G1 flies were screened for white eyes. Once we identified a *white*^-^ strain, we backcrossed the *pBac{3XP3::EYFP-attP}* markers generated previously in *D. santomea* STO CAGO 1482 to the *white*^-^ strain. The *pBac* insertion sites in these new *white*^-^ strains were then re-mapped with TagMap.

**Testing phiC31-mediated integration efficiency:** Different *attP* landing sites provide different efficiencies of integration of *attB*-containing plasmids (Bischof *et al.* 2007). We performed a preliminary screen of integration efficiency on a subset of the *attP* landing sites we generated. Pre-blastoderm embryos were co-injected with 250 ng/uL of plasmids containing *attB* sites and 250 ng/uL pBS130 (Gohl *et al.* 2011), a heat-shock inducible source of phiC31 integrase, and one hour after injection were incubated at 37°C for one hour. G0 offspring were backcrossed to the parental line and G1 offspring were screened for the relevant integration marker. We performed this screen using a heterogeneous collection of plasmids that we are integrating for other purposes. Therefore, the integration efficiencies we report are not strictly comparable between sites. Nonetheless, we were able to identify a subset of sites that provide reasonable integration efficiency and which can be made homozygous after integration of transgenes. We report these statistics for all sites that we have tested (Supplementary File: Strains and Integration Efficiencies.xlsx).

**Testing expression patterns and levels of transgenes integrated in different *attP* sites:** Different *attP* landing sites drive different levels and patterns of transgene expression (Pfeiffer *et al.* 2010). We have tested a subset of the *attP* sites in our collection for embryonic expression of an integrated *D. melanogaster even-skipped* stripe 2 enhancer (Small *et al.* 1992). A plasmid containing the *D. melanogaster eveS2-placZ* was co-injected with 250 ng/uL pBS130 into approximately ten *pBac{3XP3::EYFP-attP}* strains of each species. We isolated transgenic lines for seven *D. simulans*, four *D. mauritiana*, two *D. yakuba* strains, and four *D. santomea* strains. We performed mRNA fluorescent in situ hybridization (FISH) and imaged mid-stage 5 embryos on a Leica TCS SPE confocal microscope. (Antibody staining is less sensitive at these stages than FISH due to slow production of reporter gene protein products.) Embryos of all samples were scanned with equal laser power to allow quantitative comparisons of expression patterns between strains.

We performed staining experiments for all sites from each species in parallel; embryo collection, fixation, hybridization, image acquisition, and processing were performed side-by-side under identical conditions. Confocal exposures were identical for each series. Image series were acquired in a single day, to minimize signal loss. Sum projections of confocal stacks were assembled, embryos were scaled to match sizes, background was subtracted using a 50-pixel rolling-ball radius and fluorescence intensity was analyzed using ImageJ software (http://rsb.info.nih.gov/ij/).

**Killing *EYFP* expression from *attP* landing sites:** Expression of the *EYFP* genes associated with the *attP* sites may conflict with some potential uses of the *attP* landing sites, for example for integration of transgenes driving *GFP*-derivatives, such as *GCaMP*, in the brain. We have therefore started generating *pBac{3XP3::EYFP-attP}* strains where we have killed the *EYFP* activity using CRISPR-Cas9 mediated targeted mutagenesis. We first built a derivative of the *pCFD4-U61-U63* tandem gRNAs plasmid (Port *et al.* 2014) where we replaced the *vermillion* marker with a *3XP3::DsRed* dominant marker. The *vermillion* marker was removed by HindIII digestion of pCFD4-U61-U63 and isolation of the 5,253 bp band. The *3XP3::DsRed* cassette was amplified from a *pUC57{3XP3::DsRed}* plasmid using the following primers: 5’ TAC GAC TCA CTA TAG GGC GAA TTG GGT ACA CCA GTG AAT TCG AGC TCG GT, 5’ TTG GAT GCA GCC TCG AGA TCG ATG ATA TCA ATT ACG CCA AGC TTG CAT GC. The PCR product and vector backbone were assembled with Gibson assembly (Gibson *et al.* 2009) following http://openwetware.org/wiki/Gibson_Assembly to Generate *p{CFD4-3XP3::DsRed-BbsI}*. To remove the BbsI restriction site from *DsRed*, which conflicts with the BbsI restriction site used for cloning gRNA sequences, we digested this plasmid with NcoI and isolated the ~6kb fragment, PCR amplified this region with primers that eliminated the BbsI restriction site (Forward primer: 5’ CGG GCC CGG GAT CCA CCG GTC GCC ACC ATG GTG CGC TCC TCC AAG AAC GTC A, Reverse primer: 5’ CGC TCG GTG GAG GCC TCC CAG CCC ATG GTT TTC TTC TGC ATT ACG GGG CC), and Gibson cloned the PCR product into the plasmid backbone. This yielded plasmid *p{CFD4-3xP3::DsRed}*.

To make a plasmid for mutating *EYFP* in fly lines, we digested *p{CFD4-3xP3::DsRed}* with BbsI and gel purified the 5,913 bp fragment. A gBlocks Gene Fragment (IDT) (5’ CAA GTA CAT ATT CTG CAA GAG TAC AGT ATA TAT AGG AAA GAT ATC CGG GTG AAC TTC GGG TGG TGC AGA TGA ACT TCA GTT TTA GAG CTA GAA ATA GCA AGT TAA AAT AAG GCT AGT CCG TTA TCA ACT TG), which contained a gRNA sequence targeting EYFP that was previously validated by direct injection of gRNA was synthesized and Gibson assembled with the BbsI digested fragment of *p{CFD4-3xP3::DsRed}* to make *p{CFD4-EYFP-3xP3::DsRed}*.

This plasmid contains *attB* and can be integrated into *attP* sites. We tested this by integrating this plasmid into the *attP* site of *D. simulans* line 930. This plasmid is a potent source of gRNA targeting EYFP, which we confirmed by crossing this line to a transgenic strain carrying *nos-Cas9*. We have generated transgenic strains of *Drosophila simulans, D. mauritiana*, and *D. yakuba* carrying *nos-Cas9* (Addgene plasmid 62208 described in Port et al. 2014) and details of these lines are provided as Supplemental Data.

To knock out *EYFP* in specific strains carrying *pBac{3XP3::EYFP-attP}*, we co-injected 500 ng/uL *in vitro* transcribed Cas9 mRNA and 250 ng/uL *p{CFD4-EYFP-3XP3::DsRed}*. G0 individuals were brother-sister mated and we screened for reduction or loss of *EYFP* expression in G1 progeny. Individuals displaying reduced or no *EYFP* expression were crossed to generate strains homozygous for *EYFP*^-^.

**Data and Reagent Availability:** Plasmid *pBac{3XP3::EYFP-attP}* is available from David Stern upon request. The p{CFD4} derivative plasmids have been deposited with Addgene (plasmid IDs 86863 – 86864). All fly stocks are maintained in the Stern lab at Janelia Research Campus and all requests for fly stocks should be directed to David Stern. The raw iPCR and TagMap data are available upon request from David Stern. We continue to produce new fly strains based on the reagents described in this paper. An Excel sheet containing information about all strains in this paper and any new lines is available at http://research.janelia.org/sternlab/Strains_and_Integration_Efficiencies.xlsx. Geneious files containing genomic insertion sites for all transgenes will be updated with new strains and are available at the following sites: http://research.janelia.org/sternlab/D.simulans_mauritiana_insertions.geneious; http://research.janelia.org/sternlab/D.yakuba_santomea_insertions.geneious; http://research.janelia.org/sternlab/D.virilis_insertions.geneious. All of these files can be accessed via our lab web page at https://www.janelia.org/lab/stern-lab/tools-reagents-data.

## Results

**Generation and mapping of *pBac{3XP3*::*EYFP-attP}* strains:** We generated many strains carrying *pBac{3XP3::EYFP-attP}* and *pBac{3XP3::DsRed}* insertions, mapped these, and culled the collection to unique lines that could be maintained as homozygotes. The final collection includes 184 *D. simulans* lines, 122 *D. mauritiana* lines, 104 *D. yakuba* lines, 64 *D. santomea* lines, and nine *D. virilis* lines. Maps indicating the insertion site locations are shown in Figures 2–6 and are provided as searchable Geneious files (http://www.geneious.com/) in Supplementary Material. Details of the transgenic strains are provided in Supplementary Data: Strains and Integration Efficiencies.xlsx.

**Figure 2.**
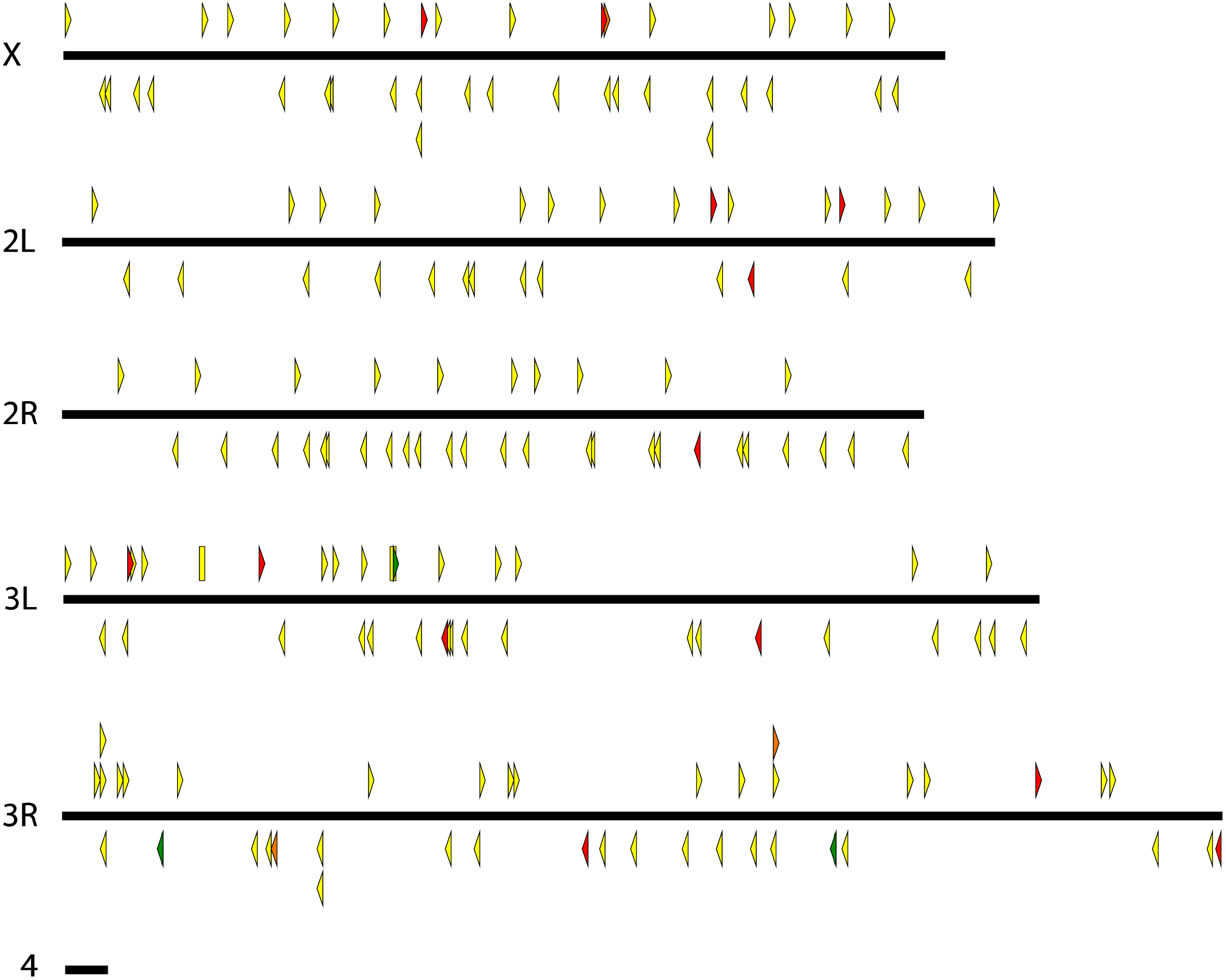
Genomic insertion sites of *pBac* transposable elements in *D. simulans*. Each triangle represents a unique *pBac* element insertion. Some strains carry multiple insertion events. Some insertion sites are present in multiple strains at least one of which contains multiple insertions. These strains were maintained to maximize the diversity of insertion sites in the collection*. pBac* insertions oriented forward are indicated above each chromosome and point to the right and reverse insertions are indicated below each chromosome and point to the left. Rectangles represent inserted elements whose orientation could not be determined. Yellow, green and red indicated elements carrying *3XP3::EYFP, 3XP3::EGFP*, and *3XP3::DsRed*, respectively.

**Figure 3.**
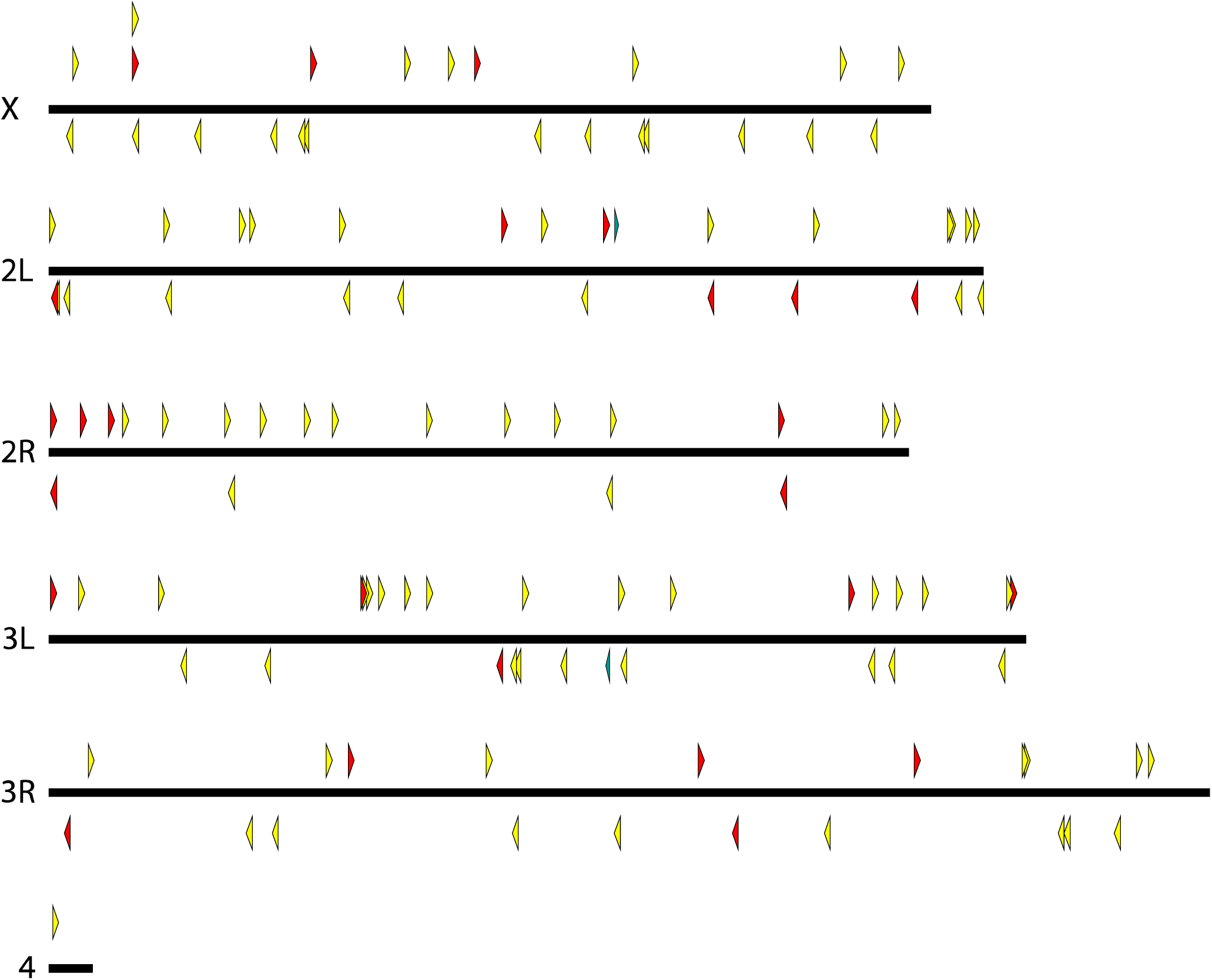
Genomic insertion sites of *pBac* transposable elements in *D. mauritiana*. All details as in Figure 2 legend.

**Figure 4.**
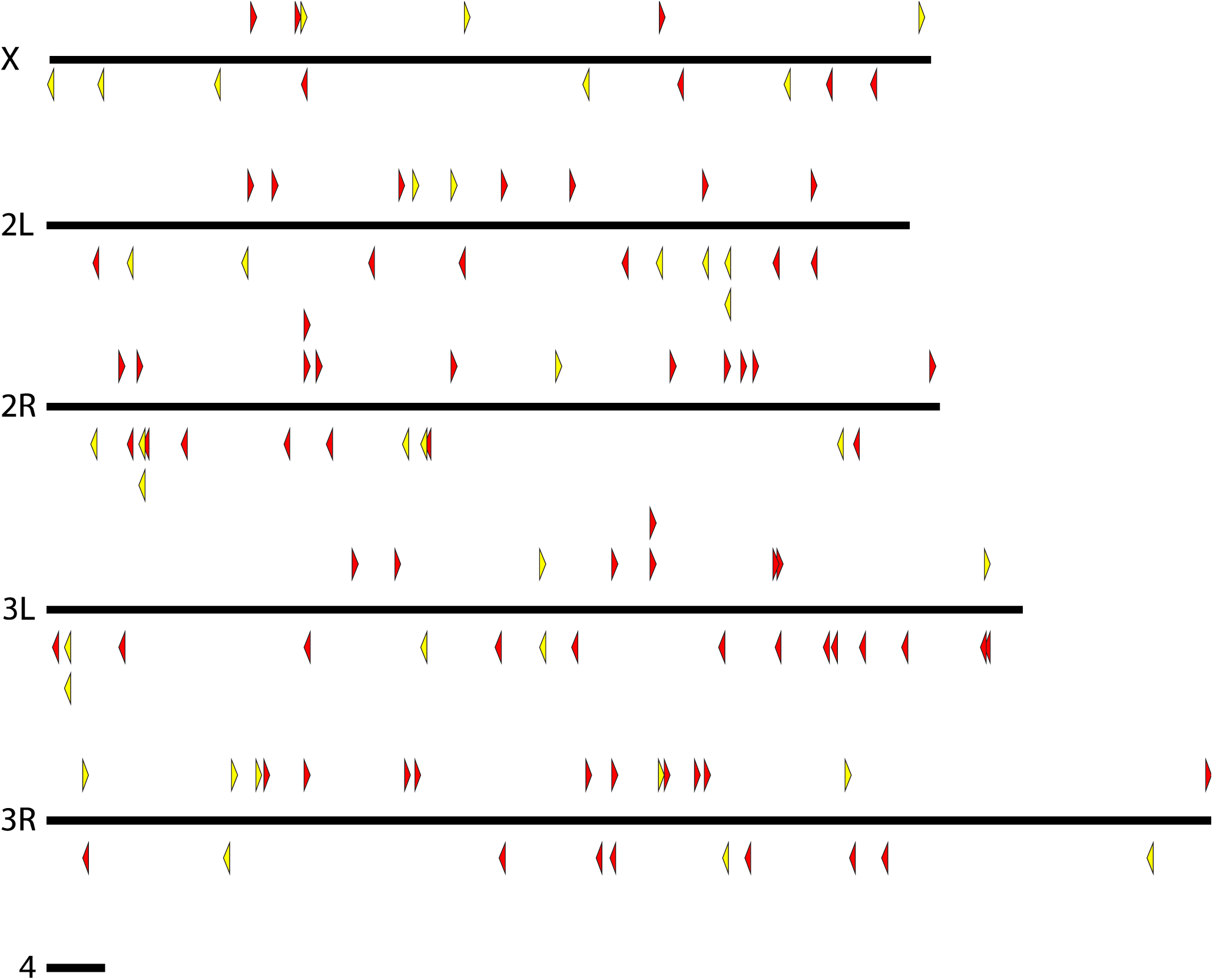
Genomic insertion sites of *pBac* transposable elements in *D. yakuba*. All details as in Figure 2 legend.

**Figure 5.**
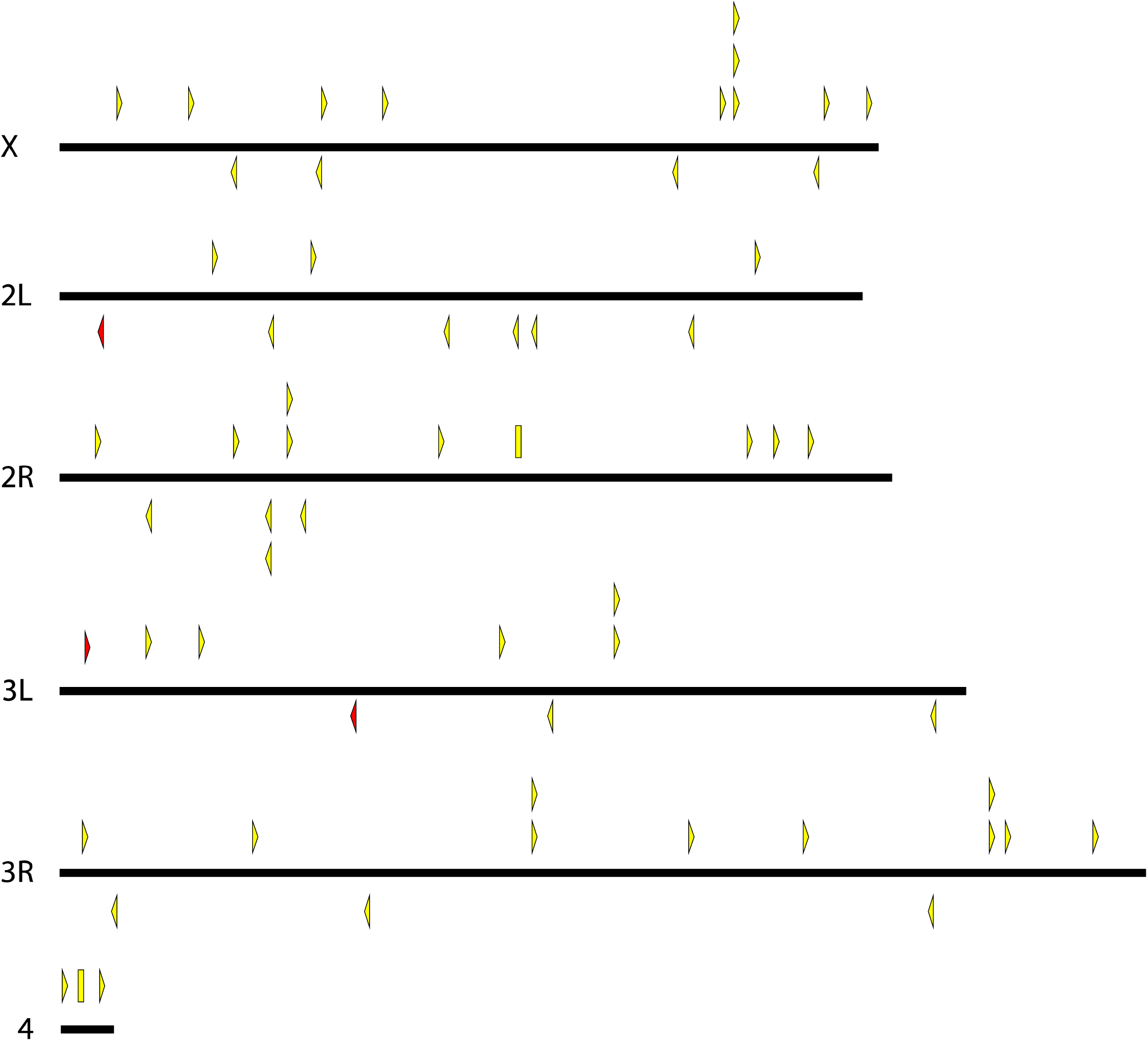
Genomic insertion sites of *pBac* transposable elements in *D. santomea*. All details as in Figure 2 legend.

**Figure 6.**
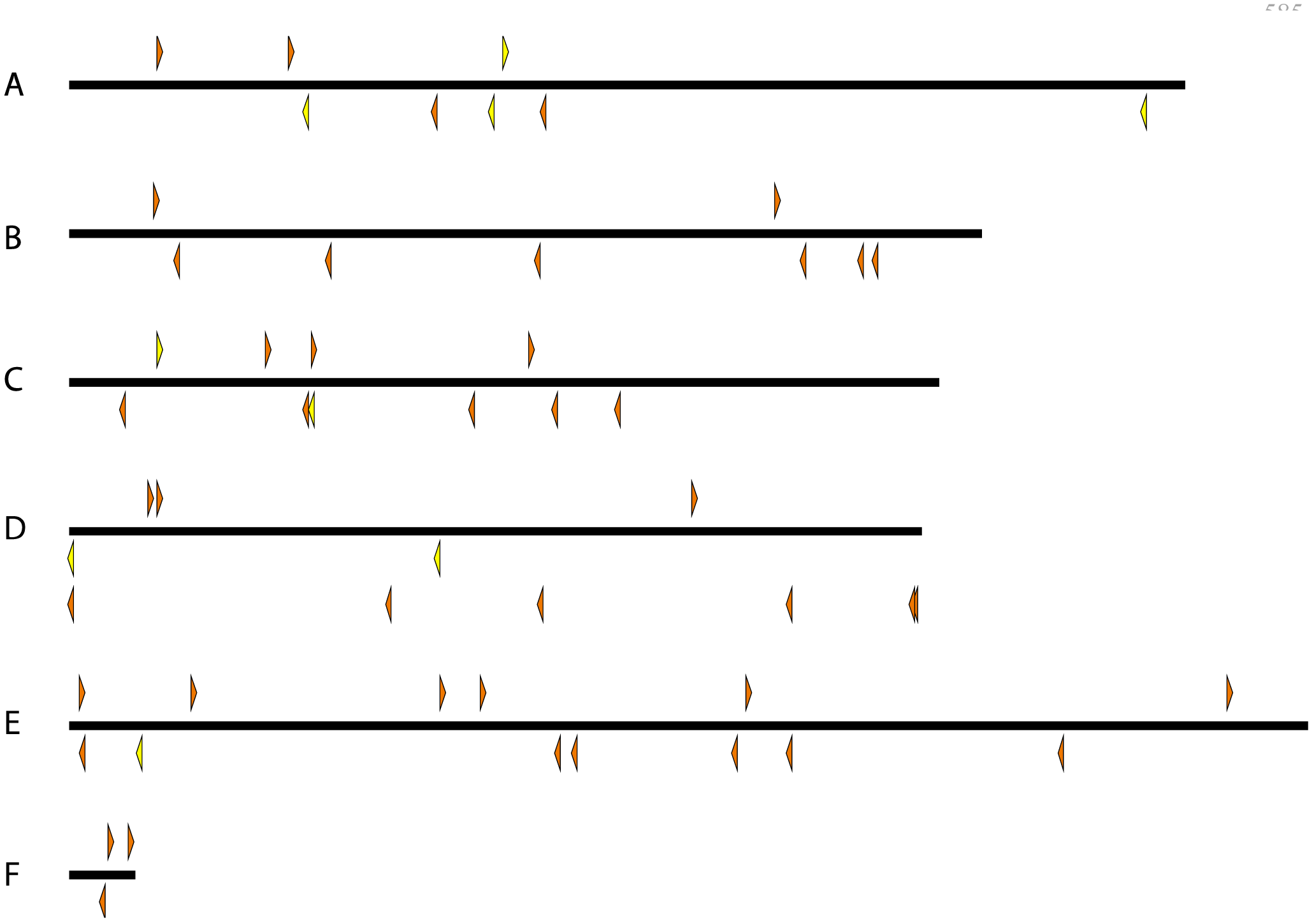
Genomic insertion sites of *pBac* transposable elements in *D. virilis*. Each triangle represents a unique *pBac*element insertion. Some strains carry multiple insertion events. *pBac* insertions oriented forward are indicated above each chromosome and point to the right and reverse insertions are indicated below each chromosome and point to the left. Yellow and orange indicate elements carrying *3XP3::EYFP* and *w*^+^, respectively.

**Mapping*pBac*transposon insertion sites in *D. virilis*:** To assist with genetic experiments in *D. virilis*, we mapped the insertion locations for all *pBac* lines generated in our lab for a previously published study (Frankel *et al.* 2012). We mapped 58 transposon insertions from 39 *pBac*(*enhancer-lacZ*) strains plus nine new *pBac{3XP3::EYFP-attP}* strains. Some strains contained multiple insertions and some insertions mapped to contigs that are not currently associated with Muller arm chromosomes. These results are shown in Figure 6 and available in a Geneious file and Supplementary Data: Strains and Integration Efficiencies.xlsx.

**Testing phiC31-mediated integration efficiency:** We tested efficiency of integration of *attB* plasmids into *attP* landing sites of multiple strains of each species. There are strong differences in integration efficiencies between landing sites. Some landing sites in *D. simulans, D.mauritiana, D. santomea* and *D. yakuba* supported integration of *attB* plasmids, although many landing sites did not support integration at reasonable frequency. Details of integration efficiencies for each line are provided in Supplementary Data: Strains and Integration Efficiencies.xlsx.

In addition, we tested nine *D. virilis* strains carrying *pBac{3XP3::EYFP-attP}* for integration of the *eveS2-placZ* plasmid, which contains an *attB* site. We screened approximately 100 fertile G0 offspring for each of these nine strains and did not recover any integrants. This is a surprising result and we do yet know whether this failure of *attB* integration is specific to these lines or reflects a general low efficiency of *attP-attB* integration in *D. virilis.*

**Testing expression patterns of transgenes integrated in different *attP* sites:** We integrated a *D. melanogaster eveS2-placZ* plasmid into multiple *attP* landing site strains of each species to examine variability in expression at different landing sites. Levels of reporter gene expression varied between strains (Figure 7). In *D. simulans, D. mauritiana*, and *D. yakuba*, we identified at least one strain that drove strong and temporal-spatially accurate levels of *eveS2* expression. However, of the four landing sites we tested in *D. santomea*, none provided strong expression of *eveS2* (Figure 7 & 8). *eveS2* transgenes often drive weak, spatially diffuse expression prior to stage 5, and all of the *D. santomea* strains displayed similar diffuse, weak expression at early stages. We also observed ectopic expression of the *eveS2* transgene in *D. santomea 2092* (Figure 8h). It is not clear if the poor expression of *eveS2* in these *D. santomea* landing sites reflects differential regulation of the *D. melanogaster eveS2* enhancer in *D. santomea* or suppression of expression caused by position effects of these specific landing sites.

**Figure 7.**
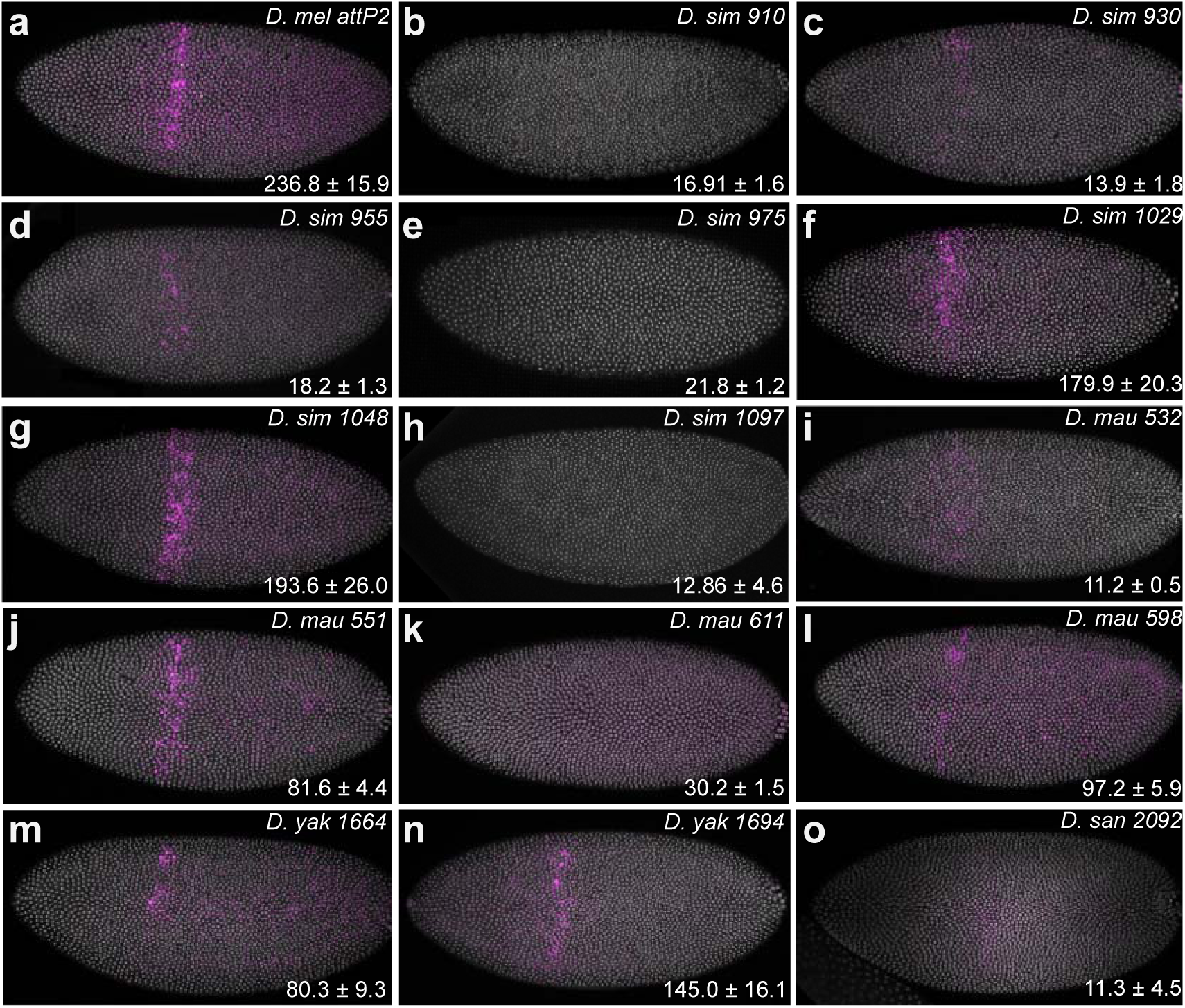
Variation in transgene expression supported by different *attP* landing sites in four species. An *eveS2* transgene driving expression in the *even-skipped stripe 2* domain of early embryos was inserted into multiple *attP* sites ofeach of four species: *D. simulans, D. mauritiana, D. yakuba*, and *D. santomea. eveS2* expression is shown in purple and DNA was counterstained with DAPI and shown in white. Expression levels in the stripe 2 domain were quantified in ten embryos of each strain and the mean ± standard deviation are reported in the bottom right corner of each panel in arbitraryunits of fluoresence intensity. (a) As a control, we stained a line containing the same plasmid inserted into the *attP2* site of *D. melanogaster*. (b-n) Seven *attP* strains of *D. simulans* (b-h), four *attP* strains of *D. mauritiana* (i-l), and two *attP* strains of *D. yakuba* (m, n) support different levels of *eveS2* expression. (n) Strain 1694 contains two *attP* landing sitesand we have not determined which landing site contains the *eveS2* transgene or whether both do. (o) None of the four *D. santomea attP* strains we tested supported high levels of spatio-temporally correct *eveS2* expression. The strain displaying the strongest expression (*2092*) is shown here.

**Figure 8.**
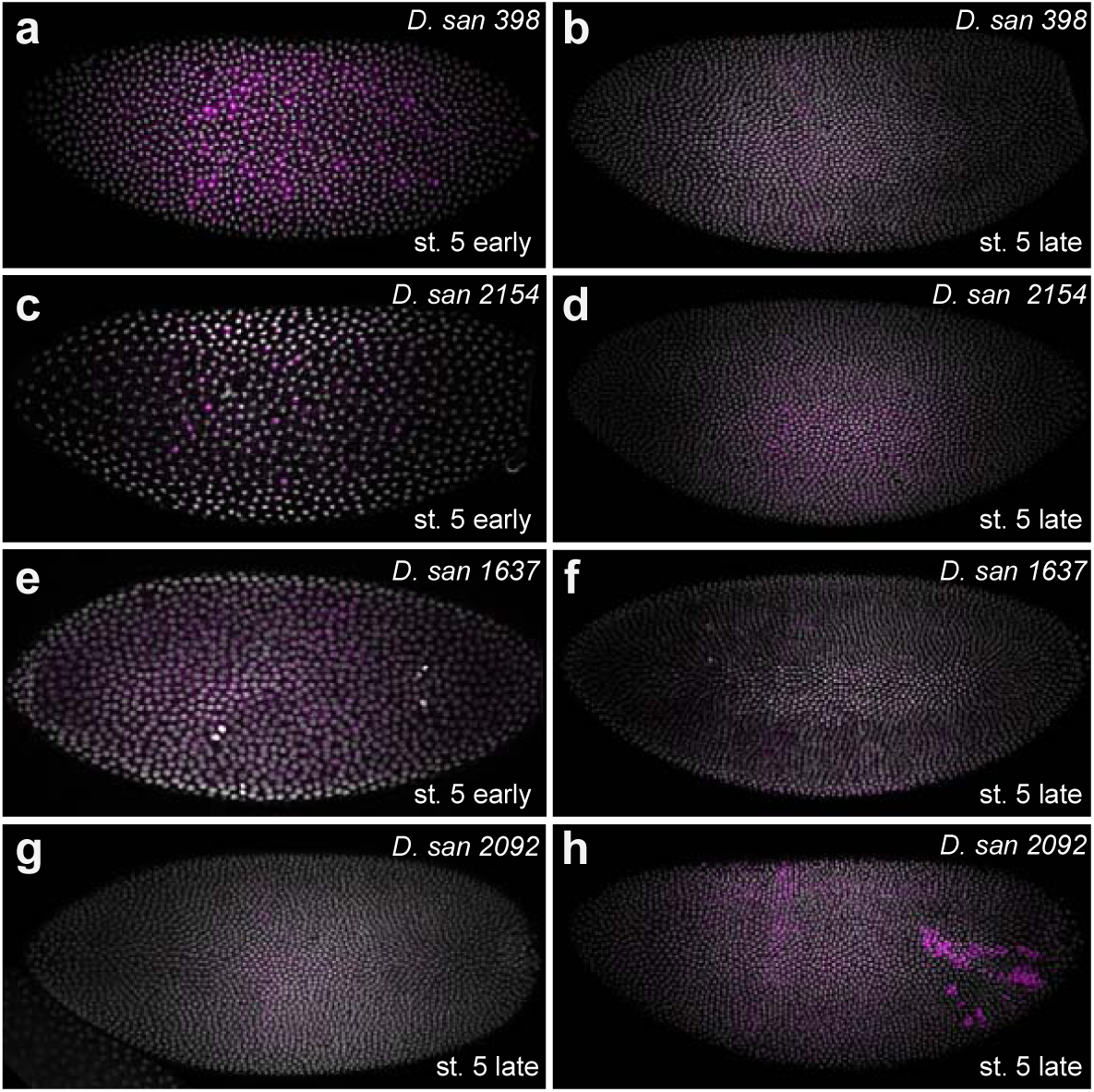
Four *D. santomea attP* landing sites do not support spatio-temporally correct *eveS2* transgene expression. (a-g) In early stage 5 embryonic stages, the lines displayed variable levels of diffuse expression, as is often observed with *eveS2* transgenes (a, c, e). However, at late stage5, none of the lines drove strong expression in the stripe 2 region (b, d, f, g). (h) Strain 2092 sometimes displayed strong ectopic expression outside of the stripe 2 domain.

**Unmarked attP landing sites:** To facilitate integration of plasmids expressing fluorescent proteins that overlap with the excitation and emission spectra of EYFP, we have generated a subset of strains in which we induced null mutations in the *EYFP* gene marking the *attP* landing sites. These strains were generated by CRISPR/Cas9-induced mutagenesis. All strains were sequenced to ensure that the mutations did not disrupt the *attP* landing site. We have so far generated two strains in *D. mauritiana*, and three strains in each of *D. santomea, D. simulans* and *D. yakuba* (Supplementary Material).

## Discussion

We have generated a collection of transgenic strains that will be useful for multiple kinds of experiments. First, the *3XP3::EYFP-attP* strains provide a collection of *attP* landing sites for each species that will facilitate transgenic assays in these species. Integration efficiencies vary widely between strains and our experiments provide some guidance to identify landing sites with the highest efficiency of integration. Second, these transgenes carry markers that will be useful for genetic mapping experiments. Several published studies have already used these reagents and illustrate the power of these strains for genetic studies (Andolfatto *et al.* 2011; Erezyilmaz and Stern 2013; Ding *et al.* 2016).

We have generated transgenic strains using these *attP* landing sites and found that they show variation in embryonic expression patterns (Figures 7 & 8). These results provide a rough guide to which strains may be useful for experiments that require low or high levels of embryonic expression. However, these results may not be predictive of transgene expression patterns at other developmental stages and in other tissues and we strongly encourage colleagues to test a variety of landing sites for their experiments and report their experiences to us. We plan to continue to maintain a database reporting on integration efficiencies and expression patterns and we will periodically update the Excel file associated with this manuscript.

This collection of reagents complements the existing resources available for studying species of the genus *Drosophila*, including the availability of multiple genome sequences (Clark *et al.* 2007) and BAC resources (Song *et al.* 2011). This resource will accelerate research on gene function in diverse *Drosophila* species and the study of evolution in the genus *Drosophila*.

## Acknowledgements

We thank Peter Andolfatto and an anonymous referee for helpful comments that improved this paper. Most fly stocks were provided by the San Diego Species Stock Center and *D. santomea* STO CAGO 1482 was kindly provided by Peter Andolfatto. We thank Ernst Wimmer for providing the original *piggyBac* reporter plasmids.

### Author Contributions

DLS conceived of the project. NF made *pBac{3XP3::EYFP-attP}*. DLS, YD, GK and JDM screened injected flies for integration events. YD, RK, AL, J-YK and SP performed iPCR experiments. JYK prepared DNA samples for TagMap. DLS performed TagMap. AL and SP sequenced the TagMap libraries. DLS, YD, GK, and JYK performed the genetics. JC performed the embryo *in situ* hybridization experiments. DLS wrote the paper.

